# A Semi-Dry, Visually Comfortable c-VEP BCI with Short Calibration, Conservative Idle-State Behavior, and a VR Proof of Concept

**DOI:** 10.64898/2026.09.29.754884

**Authors:** Frederic Dehais, Emilio Maggi, Kalou Cabrera Castillos

**Affiliations:** Fédération ENAC ISAE-SUPAERO ONERA, Université de Toulouse, France; School of Biomedical Engineering, Science and Health Systems, Drexel University, Philadelphia, PA, USA

**Keywords:** Reactive brain-computer interface, idle state, code-modulated visual evoked potential, textured visual stimuli, visual comfort, semi-dry EEG

## Abstract

**Background and Objectives:** Code-modulated visual evoked potentials (c-VEPs) provide fast and reliable control signals for reactive brain-computer interfaces (rBCIs). However, deployment beyond laboratories remains limited by visually intrusive stimulation, reliance on wet electroencephalography (EEG), and false activations during non-control periods. We introduce a usability-oriented c-VEP rBCI combining low-salience textured stimulation, slow-flashing codes, asynchronous evidence accumulation, and semi-dry EEG acquisition.

**Methods:** Thirty participants evaluated the system on a standard screen using an 11-class keypad typing task. Two identical online typing sessions were separated by a 2-min video-viewing period serving as an explicit idle condition. Feasibility was then assessed in virtual reality (VR) through a proof-of-concept shell-game task involving head movements and immersive rendering constraints. Visual comfort, peripheral distraction, and fatigue were assessed subjectively in both conditions.

**Results:** In the keypad task, the system achieved 99.8% accuracy with 33 s of calibration data and a mean decoding time of 4.3 s. No spurious commands occurred during the tested 2-min idle condition. In VR, online accuracy reached 100%, with a mean decoding time of 5.6 s. Participants reported high visual comfort and minimal peripheral distraction in both conditions, although fatigue was higher in VR.

**Conclusion:** The proposed c-VEP rBCI combines short calibration, high accuracy, comfortable stimulation, semi-dry EEG, and asynchronous decision-making. Performance during the explicit idle condition and VR implementation support the feasibility of wearable and practical c-VEP systems for self-paced interaction beyond laboratory environments. These findings encourage evaluation in longer idle periods and real-world tasks.

## 1 Introduction

Code-modulated visual evoked potentials (c-VEPs) and steady-state visual evoked potentials (SSVEPs) are among the most effective control signals for reactive brain–computer interfaces (rBCIs) [1, 2]. Their high speed and robustness make visual rBCIs particularly attractive for fast and intuitive interaction. However, despite strong laboratory performance, their translation to practical end-user applications remains limited by several persistent usability constraints [3]. First, many systems still require substantial calibration, a burden that tends to increase with the number of target classes, particularly in SSVEP-based approaches (e.g., up to 40 min in [4]). Such requirements reduce accessibility and hinder spontaneous use. Second, many high-performance implementations still depend on wet EEG, which reduces practicality outside controlled settings [5]. Finally, the stimulation itself is often highly salient, visually fatiguing, and distracting in peripheral vision, especially when multiple flickering targets are presented simultaneously [6, 7].

To address the visual-usability aspect of this problem, visual stimulation was based on the previously introduced *Stimuli for Augmented Response* (StAR) design dehais2024leveraging, inspired by the stimulation strategy implemented in the NextMind system nextmind. The present experiment used the Ricker-textured StAR configuration, in which each stimulation region was overlaid with a texture composed of small, spatially distributed Ricker patches. The StAR design is intended to exploit early visual processing mechanisms, from contrast-sensitive retinal pathways to orientation-selective cortical populations, while remaining perceptually unobtrusive, particularly in peripheral vision. Its rationale is also consistent with crowding-like visual mechanisms, whereby nearby elements become less individually distinguishable outside central vision harrison2015unifying. Previous work showed that Ricker-textured StAR stimuli can enhance visually evoked responses relative to conventional uniform flickers while improving subjective visual comfort dehais2024leveraging.

Another important challenge concerns the handling of non-control periods. In many reactive BCIs, the system is effectively required to output a decision at each decoding step; when the user is not attending to any target, looks away, or divides attention with another task, the decoder may still produce a spurious best match, leading to false activations. This issue has motivated a growing body of work on asynchronous control and idle-state detection in both SSVEP and c-VEP paradigms. In SSVEP, prior studies have proposed pseudo-key-based brain switches, explicit idle-state classifiers, and multi-stage work/idle detectors based on task-related component analysis and related methods [8, 9, 10, 11, 12]. In VEP/c-VEP systems, asynchronous non-control handling has likewise been addressed through confidence-based rejection, explicit brain-switch formulations, or calibrated thresholding strategies [13, 14, 15, 16, 17]. These studies represent important steps toward self-paced visual BCIs. However, non-control handling often requires an explicit rejection mechanism, a dedicated idle-state model, user-specific threshold optimization, or a binary brain-switch formulation. Fewer studies have examined whether idle-state behavior can emerge directly from the same evidence-accumulation process used for multi-target selection, while also addressing calibration burden, visual comfort, and hardware practicality.

To address several of these limitations jointly, we build on the Burst-code c-VEP paradigm [18], which concentrates stimulation into brief, aperiodic flashes delivered at a low rate of approximately 3 flashes/s. By time-locking stimulus energy into short two-frame flashes and spacing bursts sufficiently apart, Burst stimulation increases evoked-response amplitude relative to conventional faster m-sequence stimulation, thereby improving the signal-to-noise ratio under practical recording conditions. An additional advantage of Burst coding is that calibration is reduced to a binary discrimination between flash (’1’) and no-flash (’0’), independently of the number of simultaneously displayed targets. Online, the decoder correlates a stream of flash probabilities with predefined Burst codes and commits only once sufficient evidence has accumulated, enabling flexible decision times instead of fixed decoding windows. Combined with StAR stimuli, this property enabled, in our previous work, high-performance decoding in an 11-class keypad task using as little as one minute [19] or 33 s [20] of calibration data with dry EEG, with classification accuracy exceeding 93%. This decoding framework is also compatible with a form of implicit idle-state gating: when the user is not attending to any target, responses are expected to be dominated by “0”-like evidence, such that code correlations remain weak and no command reaches the decision threshold. However, this aspect has not yet been directly tested and is therefore a central objective of the present work.

Another important step toward practical deployment is the integration of BCIs into immersive virtual environments. Several studies have shown that performance in Virtual Reality (VR) can be comparable to, or in some cases better than, conventional screen-based interaction across multiple paradigms, including P300 [21, 22], SSVEP [23, 24], and motor imagery [25]. In contrast, c-VEP has received comparatively little attention in VR, despite its relevance for fast reactive control. To date, the closest demonstration is a mixed-reality c-VEP speller evaluated in a head-mounted display under largely static viewing conditions, in which a two-dimensional interface was overlaid on a controlled passthrough background and participants remained seated to limit motion artifacts [26]. Although this work constitutes an important first step toward head-mounted c-VEP interaction, it does not address more embodied scenarios in which targets are embedded in a three-dimensional environment and interaction naturally involves head motion.

In the present study, we extend this line of work through two complementary demonstrations rather than a direct comparison between screen-based and VR interaction. First, we revisit the screen-based 11-class StAR–Burst keypad paradigm introduced in [20] to explicitly assess idle-state behavior during a non-control period. Specifically, we tested whether the same decoder used for multi-target selection would remain conservative during a video-viewing idle condition, without requiring a dedicated idle-state model. Second, we implemented the same stimulation and decoding principles in a virtual-reality shell-game task, where textured c-VEP stimuli were rendered on three-dimensional objects and updated with the participant’s head motion. This experiment was intended as a proof of concept for deploying the fast-calibrated, comfort-oriented StAR– Burst framework in a head-mounted, object-based interaction context, where viewpoint changes, rendering constraints, motion-related artifacts, object-locked stimulation, and headset limitations may challenge c-VEP decoding. Participants always completed the screen-based experiment before the VR experiment; accordingly, any statistical comparisons between the two implementations were used only to characterize differences in signal properties, subjective ratings, and timing, not to establish controlled superiority or equivalence between PC and VR conditions. Both experiments were conducted using semi-dry electrodes, which offer a practical compromise between dry and wet systems by combining rapid application with improved electrode–skin contact stability, reduced sensitivity to motion-related artifacts, and enhanced wearability [27, 28, 29].

## 2 Keypad idle-state experiment: Materials and Methods

### 2.1 Participants

The study involved 30 healthy participants (10 women; mean age = 28.4 *±* 7.7 years), primarily staff and students at ISAE-Supaero. The protocol was approved by the University of Toulouse ethics committee (CER approval number 2025_1012) and complied with the Declaration of Helsinki. All participants provided written informed consent prior to participation. Exclusion criteria included a history of neurological disorders or the use of psychoactive medication at the time of testing. All participants had normal or corrected-to-normal vision and completed a video-game proficiency questionnaire.

### 2.2 Keypad PC-based BCI

The keypad-based BCI interface was presented on a 24.5-inch LCD monitor (Iiyama Gold Phoenix G-Master GB2590HSU-B1; 1920 *×* 1080; 400 cd*/*m^2^) at a refresh rate of 60 Hz. The interface consisted of an 11-class keypad inspired by traditional mobile-phone layouts, with selectable keys corresponding to digits 0–9 and the ‘#’ character. Participants were instructed to type predefined four-character sequences by overtly attending to the desired key. The current target sequence was displayed in the lower-left corner of the screen.

The BCI also provided *pre-decision visual feedback*: as decoder confidence increased for a given target (e.g., a digit or the ‘#’ key), the corresponding key was progressively enlarged as implemented in [20]. This cue allowed participants to anticipate the forthcoming decision and verify that the system was converging toward the intended target, thereby externalizing the evolving decision state before commitment and, as shown in our previous study, enhancing the perceived sense of control. After each selection, the chosen key was highlighted in green if correct or red if incorrect, providing immediate performance feedback. A video illustrating the PC condition is available here: PC condition video.

### 2.3 StAR stimuli

Visual stimulation was based on our previously introduced *Stimuli for Augmented Response* (StAR) design [19], adapted from [30]. Here, we used the *Ricker* -based version of StAR, in which each of the 11 keypad targets was overlaid with an identical texture composed of 75 small patches randomly distributed within the stimulation area (see the upper-left panel of Fig. 2). Each patch was generated from an adapted Ricker wavelet and measured 40 px *×* 20 px, corresponding to a 2:1 width-to-height ratio. Using the same texture across all targets was intended to reduce local visual distinctiveness between neighboring items, thereby increasing peripheral transparency. This effect likely relies on crowding-like mechanisms, whereby surrounding stimuli become less individually salient when viewed outside the fovea [31]. Compared with conventional high-contrast black–white alternating flickers, this texture-based stimulation was selected to improve peripheral transparency and visual comfort while amplifying evoked responses by a factor of approximately 2–3 [19]. Stimuli were presented on a medium-gray background corresponding to 50% screen luminance and modulated at a depth of 70%, a setting chosen to preserve classification performance while limiting visual salience [20].

### 2.4 Burst codes

Each of the 11 targets was assigned a unique Burst code [18]. Burst onsets were derived from a common *master code* and distributed pseudo-randomly across the 11 target codes, following the approach introduced in our previous work [18]. The *master code* was 2.2 s long on a 60 Hz display, corresponding to 132 frames, with a minimum inter-burst interval of 32 ms. It was then partitioned to generate the 11 codes used in the keypad experiment (Fig. 1).

**Figure 1:**
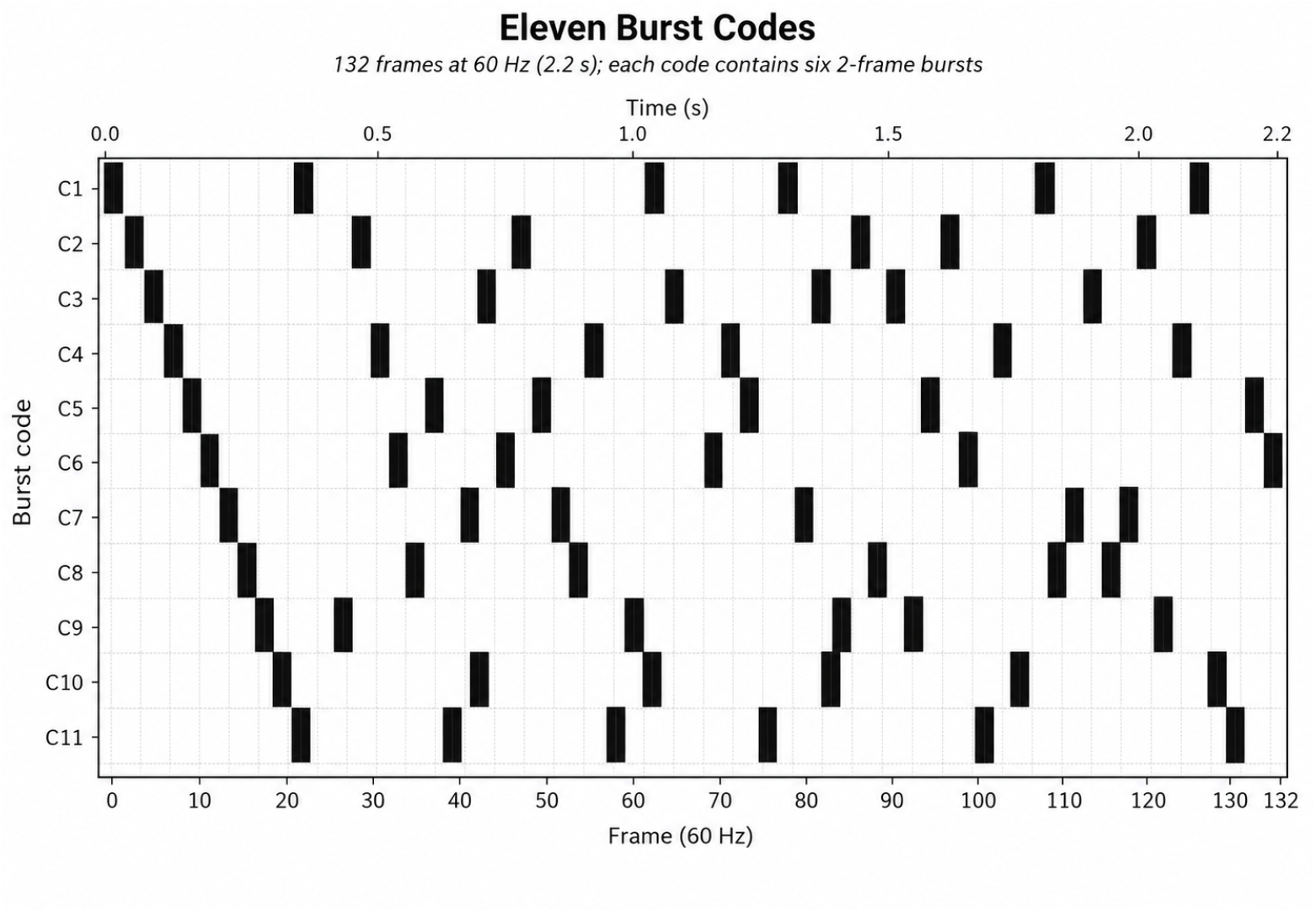
Temporal structure of the 11 Burst codes used in the keypad experiment. Each row corresponds to one target-specific code (C1–C11), defined over 132 frames at 60 Hz, corresponding to a 2.2 s code cycle. Black bars indicate two-frame burst events. Each code contains six bursts, with target identity encoded by the temporal spacing of burst onsets rather than by the number of flashes.

Each code contained six burst onsets, such that differences between codes arose from the temporal spacing of these onsets rather than from their total number. Across codes, inter-burst intervals ranged from 8 frames (132 ms) to 40 frames (660 ms). These constraints were chosen to minimize overlap between successive evoked responses while preserving sufficient temporal variability for robust code discrimination.

### 2.5 Experimental protocol

Participants were seated comfortably in front of the screen. After providing informed consent, they were fitted with an 8-channel semi-dry EEG system (MBrainTrain; 500 Hz). Electrodes were positioned over occipital and parieto-occipital regions (PO7, O1, Oz, O2, PO8, PO3, POz, and PO4). Figure 2 provides a schematic overview of the experimental protocol.

**Figure 2:**
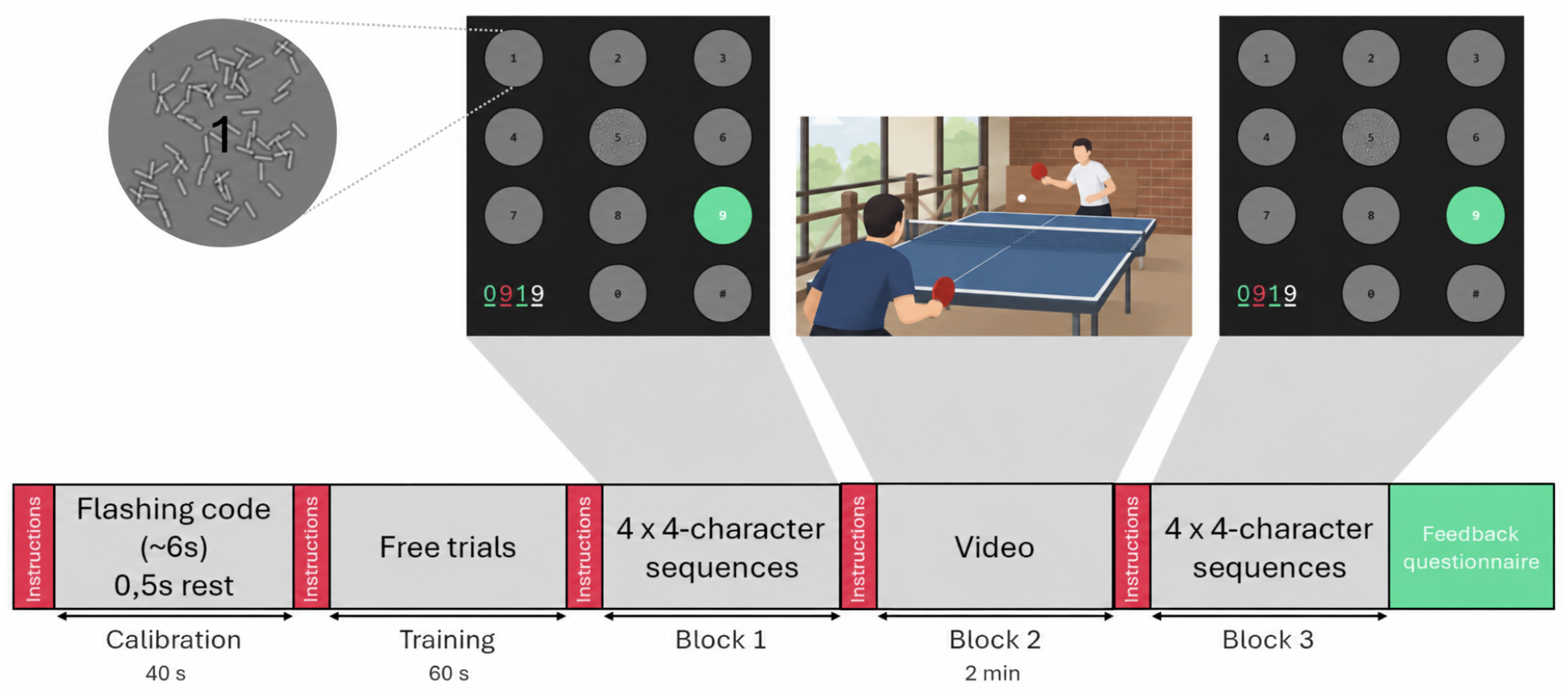
Overview of the screen-based keypad protocol. The upper panels show the keypad interface, the video-viewing period, and an example of the StAR texture-based visual stimulus used for the keypad items, illustrated here for target 1. The experiment began with a 40 s calibration phase, yielding 33 s of usable EEG data, during which participants viewed the flashing codes separated by 0.5 s rest periods. After a short training block, participants completed a first self-paced spelling block requiring four predefined 4-character sequences, corresponding to 16 selections. Idle-state behavior was then assessed during a 2-minute video-viewing period, during which the BCI pipeline remained active while no intentional control was required. Participants subsequently completed a second self-paced spelling block with four additional 4-character sequences, yielding 32 selections in total. A video demonstrating the BCI in the PC condition is available here: [PC condition video](https://nextcloud.isae.fr/index.php/s/aSGf2ZTtfi6QGGn).

Each session began with a passive calibration phase used to train the classification pipeline to detect VEP responses to flash events. During calibration, participants fixated a central circle while Burst flickers were presented in five blocks, each comprising three code presentations (2.2 s per presentation). Including cueing and rest periods, calibration lasted 40 s in total and yielded 33 s of usable EEG data. After calibration, participants were then allowed to perform free trials for up to 1 min to familiarize themselves with the upcoming task.

Participants then completed an online typing task using the BCI keypad interface. The task consisted of two spelling blocks separated by an idle-state period. In each spelling block, participants entered four predefined four-character sequences, yielding a total of 32 target selections across the experiment.

Idle-state reliability was assessed during the intermediate pause, during which participants watched a 2-minute ping-pong video while no intentional control was required. System output was monitored throughout this period to quantify potential spurious activations in the absence of voluntary BCI use.

After completion of the experiment, participants provided subjective ratings on three 10-point visual semantic differential scales assessing peripheral distraction, visual comfort, and visual fatigue. They also completed a Computer Vision Syndrome (CVS) questionnaire to verify that the task did not induce adverse visual effects.

### 2.6 Decoding pipeline

The online BCI decoding pipeline for the PC condition was implemented in *Timeflux* [32], an open-source Python framework for real-time acquisition, processing, and classification of time-series data. The user interface was implemented as a web application (HTML/CSS/JavaScript) running in a standard browser and connected to the *Timeflux* processing graph. Signal processing and classification were implemented in Python using the *imblearn* and *pyriemann* libraries. The overall software architecture follows our previous implementation [19], with minor adaptations for the present PC configuration.

EEG streams were linked to *Timeflux* via LabStreamingLayer (LSL), then preprocessed using an IIR band-pass filter (1–25 Hz) and a band-stop filter (48–52 Hz) to attenuate slow drifts and power-line noise. The filtered signal was segmented into 300 ms epochs time-locked to the display refresh cycle. Epoch timestamps were additionally shifted to compensate for an estimated 80 ms end-to-end latency in stimulus delivery and acquisition.

For each epoch, spatial information was summarized by estimating a covariance matrix across channels [33]. Prior to covariance estimation, stimulus-locked activity was enhanced and dimensionality reduced using xDAWN spatial filtering [34], retaining four spatial filters. Feature extraction was implemented with *pyriemann* (*XdawnCovariances*), which computes Riemannian “super-epochs” by concatenating each spatially filtered epoch with the corresponding average evoked response. The resulting covariance matrices were mapped to the tangent space of the Riemannian manifold using the mean covariance as the reference point. Tangent-space features were then downsampled and classified with a linear discriminant analysis (LDA) model, yielding an epoch-wise probability that a burst-evoked response was present.

Because the classifier operated at approximately 60 Hz, a single evoked response could be detected over multiple consecutive frames. To prevent duplicate detections, probabilities were binarized (threshold = 0.5) and a truncation rule was applied such that, for any run of consecutive detections, only the first frame was retained.

Calibration was used to train a binary classifier discriminating epochs elicited by a flash (‘1’) from epochs associated with a no-flash event (‘0’). The calibration session lasted 40 s in total and yielded 33 s of usable EEG after exclusion of non-stationary segments. EEG was segmented into stimulus-locked epochs; epochs following flash onsets were labeled as class 1, whereas epochs following no-flash events were labeled as class 0. During online operation, the trained classifier produced, for each incoming epoch, a probability of class 1 (flash) versus class 0 (no-flash).

Multi-class selection was then achieved by code reconstruction: the recent stream of 0/1 decoded bits was continuously matched to each target’s predefined Burst code. At each video frame, the correlation between the decoded bits stream and each class-specific code was computed, and the best-matching candidate was identified. To convert these continuously updated correlations into a robust decision with flexible timing, we used a consecutive-winner strategy with a momentum–decay dynamic [35]. The leading candidate accumulated momentum with an increment that increased as it remained in the lead, while all other candidates decayed by the same increment, bounded by a minimum floor.

The number of consecutive frames during which the same class remained the most probable was tracked and normalized by a predefined threshold of 50 frames. The updated momentum for the best candidate *I*_*N*_ and all other classes *i* at frame *N* was defined as

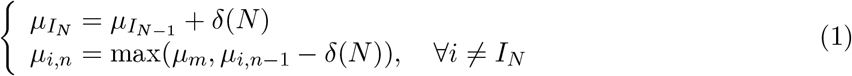

where *δ*_*i*_ denotes the momentum increment for target *i* at frame N, and *µ*_*m*_ = *−*4 defines the minimum floor preventing excessive decay. In this experiment we used the function *δ*(*N*) = 2^*N/M*^ *−*2^*N −*1*/M*^, for any frame *N* and a max frame *M* = 50. Once any class reached the maximum threshold *µ*_*M*_ = 1 for a minimum of 50 frames, it was confirmed as the selected output.

Threshold values were tuned in pretests to balance speed and robustness to transient misclassifications using an off-line data collection with 20 participants that is not presented here (classifier probability buffer: 30–50 frames; momentum starting value: *−*2; momentum floor: *−*4; minimum persistence at threshold before decision: 50 frames). Overall, this strategy ensured that selections were emitted only after sufficiently strong and temporally consistent evidence had accumulated, thereby limiting the impact of transient errors while maintaining responsiveness.

### 2.7 Subjective metrics

After each typing block, participants completed a short subjective questionnaire composed of three 10-point semantic differential scales assessing perceived visual comfort, mental fatigue, and peripheral transparency (i.e., the extent to which neighbouring flickers were distracting in peripheral vision). Because the mental fatigue scale was formulated with higher scores representing lower fatigue, responses were reverse-scored before analysis using *s*^*′*^_*c*_ = 10 *− s*_*c*_ (scale range: 0–10). After inversion, higher scores indicated greater mental fatigue.

### 2.8 BCI performance analysis

Objective BCI performance was evaluated during the online keypad task using typing accuracy and decoding time as primary metrics. Accuracy was defined as the proportion of correctly selected targets, whereas decoding time corresponded to the time elapsed between trial onset and confirmed target selection.

In addition to these behavioral performance measures, we performed an EEG analysis using MNE (v1.11.0). Continuous EEG data were band-pass filtered between 1 and 25 Hz using a third-order IIR filter with cutoff frequencies of 0.05 and 25.05 Hz. A third-order IIR band-stop filter was also applied between 48 and 52 Hz to attenuate power-line noise. The onset of each visual stimulus was defined as the time point at which the corresponding code sequence transitioned from a dark to a bright state. Continuous EEG data were then segmented into stimulus-locked epochs ranging from -0.1 to 0.5 s relative to visual-stimulation onset. For each participant, visual evoked potentials (VEPs) were obtained by averaging these epochs across trials during the keypad task at electrode Oz, where the VEP response was most pronounced. This response is shown as the blue ERP trace in Fig. 6.

To characterize system behavior during non-control periods, we complemented the online performance metrics with two analyses comparing task and idle states. The first analysis quantified the strength of stimulus-locked EEG responses using a signal-to-noise ratio (SNR). Importantly, this analysis was based on the same stimulus-locked epochs used to compute the ERPs described above, ensuring that both analyses referred to the same visual-stimulation events and preprocessing steps. This SNR should be interpreted as an aggregate evoked-response measure rather than as a single-trial estimate of decoder reliability. For each participant and condition, the SNR was computed from the averaged evoked response as the decibel ratio between the post-stimulus RMS amplitude in the 0–500 ms window (*RMS*_*Sig*_) and the pre-stimulus baseline RMS amplitude in the -100–0 ms window (*RMS*_*Base*_):

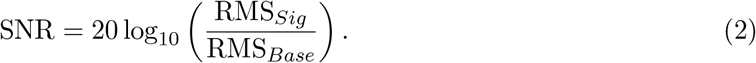

This analysis was performed separately for task periods, during which participants intentionally attended to BCI targets, and idle periods, during which they watched a video while the BCI pipeline remained active. To ensure balanced comparisons between conditions, epochs were randomly subsampled to match the minimum number of available epochs across participants and conditions, corresponding to 137 epochs per participant. This comparison was intended to assess whether stimulus-locked responses were reduced during idle periods, rather than to provide a direct estimate of the false-positive rate.

The second analysis quantified how close the decoder evidence came to the decision threshold during task and idle states. The online decoder continuously correlated the recent stream of flash probabilities with each target-specific Burst code. At each time point, the highest correlation across target codes therefore reflected the closest approach of the system to a possible decision. During intentional control, this value was expected to increase as evidence accumulated for the attended target. During idle periods, by contrast, the absence of target-related responses was expected to keep target-code correlations lower, thereby reducing the likelihood that the evidence accumulation process would reach the decision threshold.

For each participant, trial, and time point, we extracted the maximum correlation across all target codes. These values were then summarized as a distance from the decision threshold. In practice, this metric corresponds to one minus the maximum target-code correlation reached during a trial, averaged across trials. The correlation-based safety-margin metric for one participant was defined as:

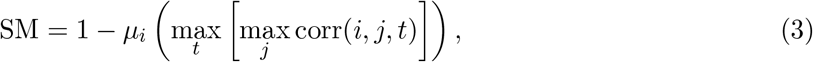

where corr(*i, j, t*) denotes the Pearson correlation between the decoded bit stream and the Burst code of target *j*, trial *i*, and time point *t*. The operator *µ*_*i*_(*·*) denotes averaging across trials.

This metric provides a continuous estimate of how far the strongest decoder evidence remains from the decision threshold, which was set to 1 in the momentum-based decision rule. Larger values indicate a greater safety margin, meaning that decoder activity remains farther from the decision boundary and is therefore less likely to produce a false activation under the tested conditions. Conversely, smaller values indicate that decoder output approaches the threshold more closely. Comparing this metric between task and idle periods thus provides a complementary assessment of whether idle-state activity remains separated from intentional BCI control.

### 2.9 Results

Normality of the data was assessed using the Shapiro-Wilk test with alpha = 0.05. No substantial deviations from normality were observed for the metrics, except for subjective ratings, which were based on Likert-type scales.

#### 2.9.1 Subjective results

Subjective ratings remained consistently favorable as participants reported high comfort (median = 8.0, IQR = 2.0), good peripheral transparency (median = 8.0, IQR = 1.0), and low mental fatigue (median = 1.0, IQR = 1.0).

#### 2.9.2 BCI accuracy and decoding time

Online control in the keypad task reached 99.8% accuracy (only one participant experienced 2 incorrect selections). The mean decoding time was 4.3 s (SD = 1.6 s) with 95% of data below 8.6 s.

#### 2.9.3 Idle-state

No false selections were observed over 60 participant-minutes of idle operation, corresponding to an observed false activation rate of 0 events/min during the video-viewing phase. During this period, participants were not intentionally attending to any BCI target, while the decoding pipeline remained active. This result provides evidence of idle-state behavior under the present experimental conditions, although it should not be interpreted as demonstrating a strictly zero false-positive rate. The momentum traces shown in Fig. 3 illustrate this behavior: during intentional control, decoder evidence progressively accumulated toward the decision threshold, whereas during the idle-state period, evidence remained below threshold in the absence of intentional BCI use.

**Figure 3:**
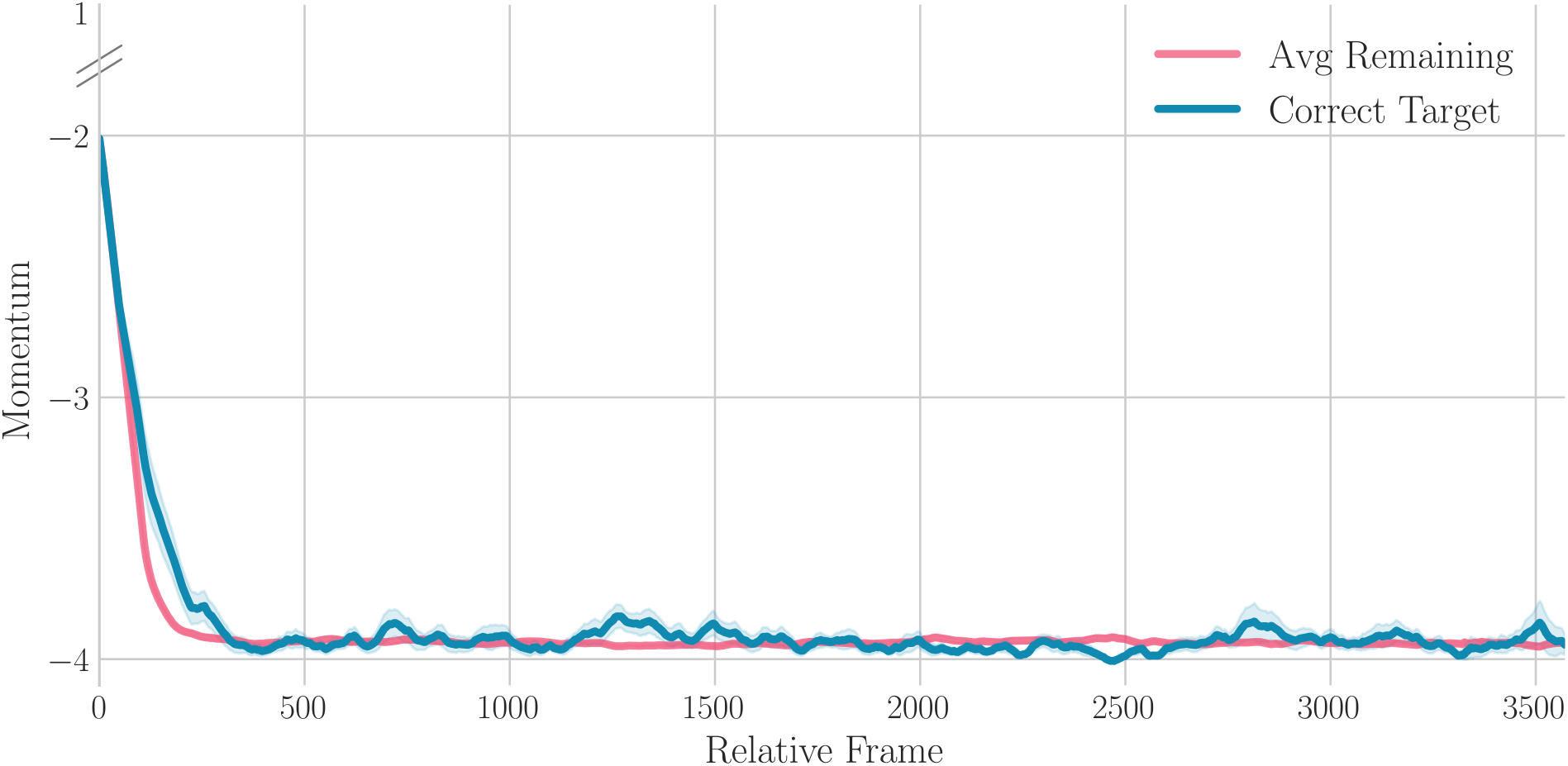
Momentum dynamics during the idle-state period. Decoder evidence remained below the decision threshold throughout the non-control period, indicating that no command was emitted while participants were watching the video and not intentionally attending to any BCI target.

As shown in Fig. 4, the Signal-to-Noise Ratio (SNR) was significantly higher during task than during idle periods (paired *t*-test, *t*(29) = 9.17, *p <* .001, *d*_*z*_ = 1.67). This large effect indicates that stimulus-locked responses were substantially reduced when participants were not engaged in intentional BCI control.

**Figure 4:**
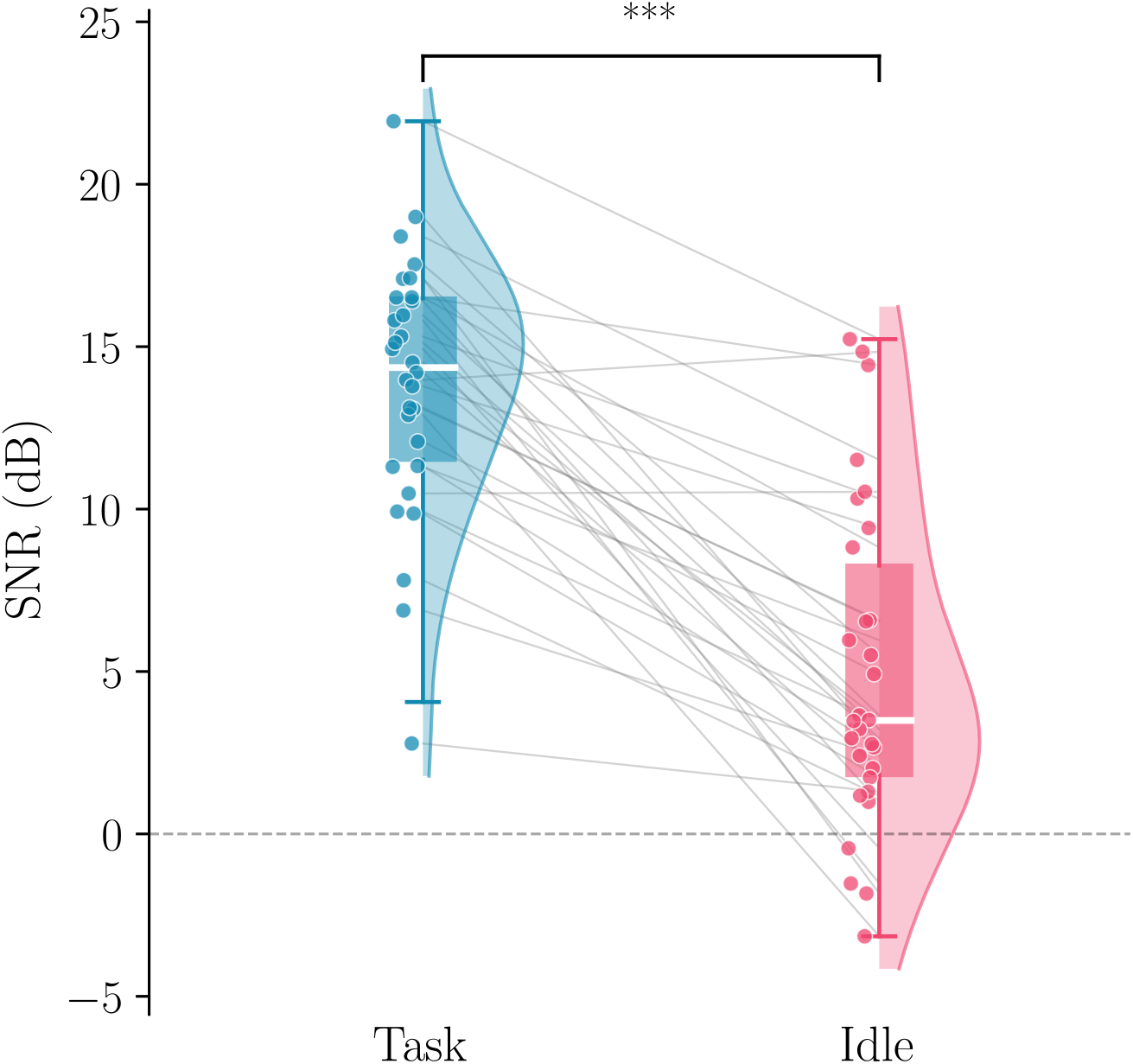
Stimulus-locked EEG signal-to-noise ratio (SNR) during task and idle periods. SNR was computed from *N* = 137 randomly subsampled epochs per participant and condition. Pairwise comparisons were performed using paired *t*-tests; three asterisks denote *p <* .001.

Likewise, the correlation-based safety-margin metric was significantly larger during idle periods than during task periods (paired *t*-test, *t*(29) = *−*78.04, *p <* .001, *d*_*z*_ = *−*14.25), suggesting that decoder evidence remained farther from the decision threshold during non-control periods. This separation is illustrated in Fig. 5, where idle periods show a larger safety margin than intentional-control periods.

**Figure 5:**
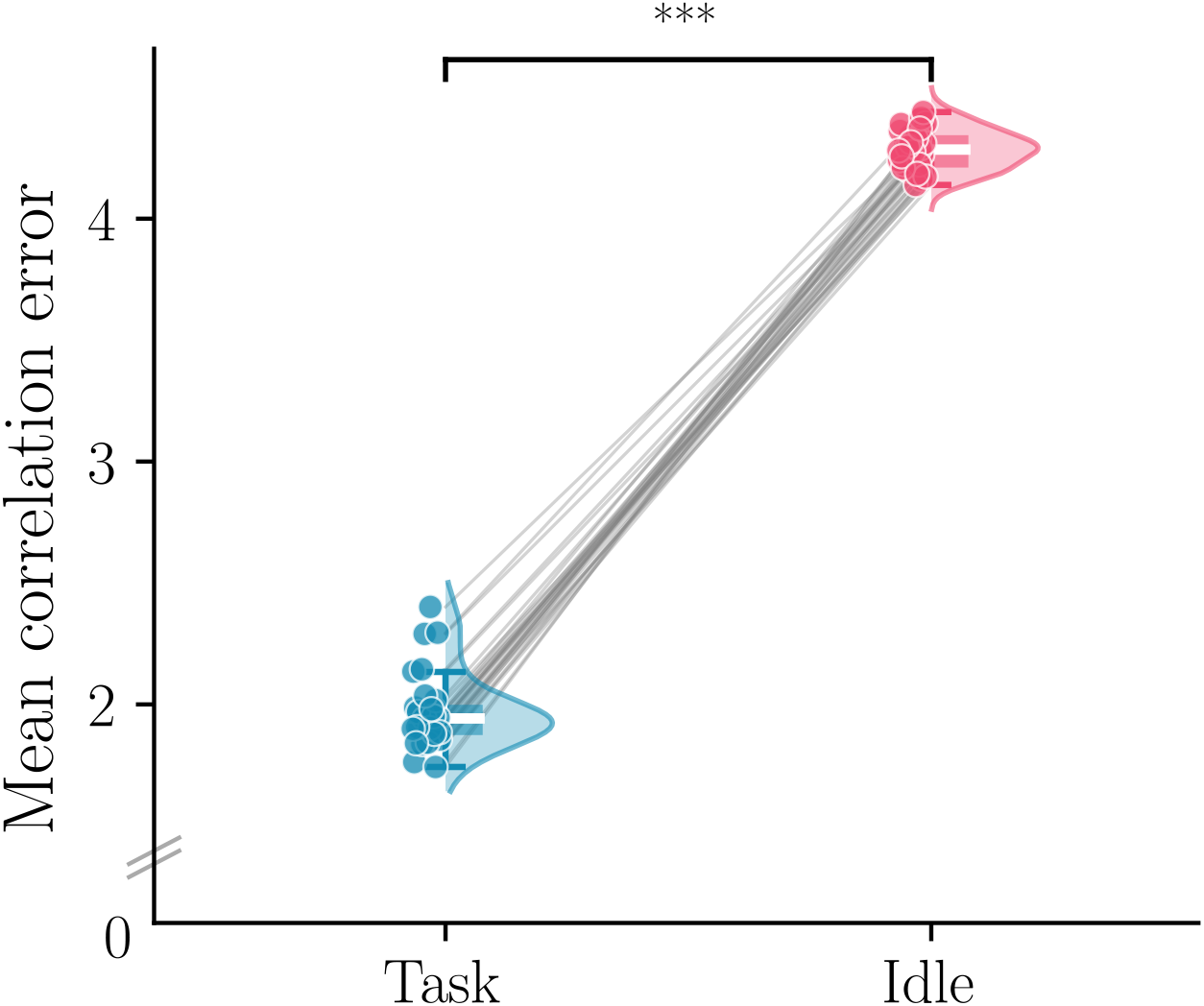
Correlation-based safety margin during task and idle periods. The safety margin reflects the distance between the maximum target-code correlation and the decision threshold, with larger values indicating that decoder evidence remained farther from the decision boundary. Pairwise comparisons were performed using paired *t*-tests; three asterisks denote *p <* .001.

### 2.10 Discussion of Study 1: screen-based keypad task

This first experiment was designed to revisit our screen-based 11-class StAR–Burst keypad paradigm while directly addressing a practical question that had remained open in our previous work: whether the Burst decoding pipeline would provide evidence of idle-state behavior without requiring a dedicated idle-state model. Specifically, we tested whether the same decoder used for multi-target selection would remain conservative during an explicit non-control period, in which participants were visually engaged in a video-viewing task while no intentional BCI control was required. The results were highly encouraging. Online control reached 99.8% accuracy, with only two incorrect selections for a single participant and perfect performance for all remaining participants, after only 33 s of usable calibration data and with a mean decoding time of 4.3 s. These findings indicate that reliable c-VEP control can be achieved with a practical semi-dry EEG configuration and a very limited calibration burden.

Compared with our previous screen-based implementation using *dry* electrodes, which achieved 93.3% mean accuracy in the same 11-class keypad task [20], the present results are compatible with the hypothesis that semi-dry contact improved signal stability [27, 28, 29]. This interpretation is consistent with the larger stimulus-locked responses observed during the calibration phase (approximately *−*5.5 *µ*V followed by +5.7 *µ*V, see Figure 6) compared with our prior dry-electrode recordings (about *−*3*µ*V followed by +1*µ*V in [19]). Because the present decoder reconstructs target identity from the inferred flash/no-flash stream, even modest improvements in elementary 0/1 discrimination are expected to propagate into more reliable multi-class code identification. Subjective ratings were also in line with our earlier reports that StAR stimulation is visually comfortable and only minimally distracting in peripheral vision [19, 20], suggesting that this gain in performance was not achieved at the expense of visual usability. This is an important usability feature for end users, as rBCIs typically rely on flashing stimuli that can be visually tiring and distracting, thereby making the interface difficult to focus on and operate. This property likely reflects the intrinsic design of the small Ricker patches, which were specifically developed to provide low-salience visual stimulation while preserving reliable c-VEP responses [30].

**Figure 6:**
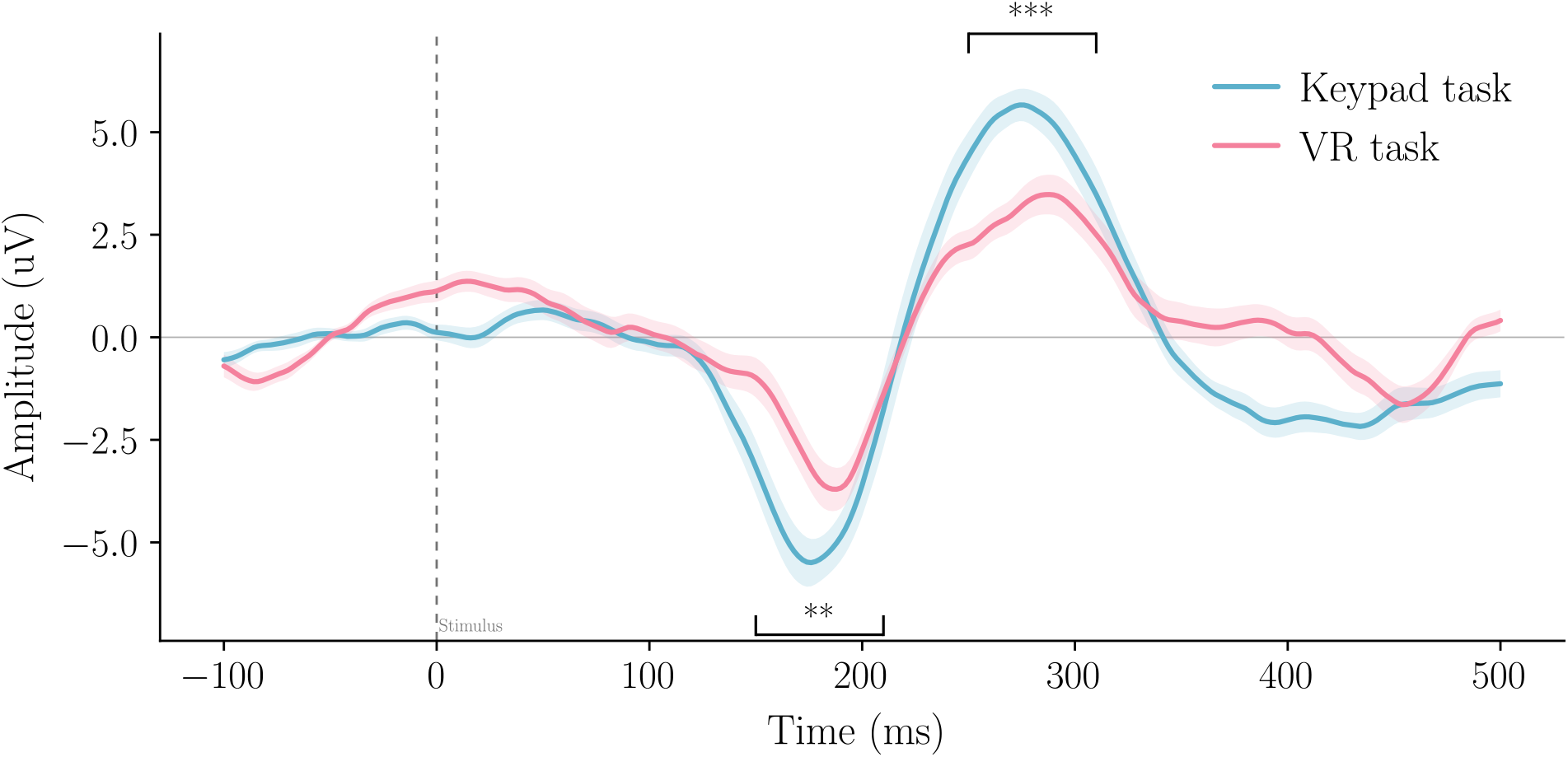
Mean evoked responses to flashes during the keypad and VR calibration phases at electrode Oz. Shaded regions indicate the standard error of the mean across participants. Paired *t*-tests were performed between conditions in the N2 time window (150–210 ms) and the P3 time window (250–310 ms). Two asterisks denote *p <* .01 and three asterisks denote *p <* .001.

A second important result concerns the handling of non-control periods. In the present experiment, idle-state behavior was tested explicitly rather than inferred indirectly: participants watched a video for approximately 2 min while the BCI pipeline remained active and no intentional control was required. This video-viewing period provided a more realistic non-control condition than simple eyes-open rest, as participants were visually engaged while the decoder continued to operate. Across 60 participant-minutes of idle operation, no spurious activations were observed. This corresponds to an observed false-activation rate of 0 events/min under the present experimental conditions, but it should not be interpreted as evidence of a strictly zero false-positive rate.

Importantly, this behavior was obtained without training a separate idle-state classifier and without introducing a dedicated calibration phase for non-control periods. Instead, idle-state handling emerged from the same Burst-code reconstruction and evidence-accumulation process used for intentional target selection. During intentional control, stimulus-locked responses and target-code correlations provide consistent evidence for the attended item, allowing momentum to accumulate toward the decision threshold. During idle periods, by contrast, the absence of consistent correlation between decoded bit stream and Burst codes is expected to keep correlations with all candidate codes low, preventing the accumulation process from reaching threshold.

The complementary EEG and decoder-level analyses support this interpretation. Stimulus-locked SNR was significantly lower during idle periods than during task periods, indicating that visual responses to the flickering codes were strongly reduced when participants were not intentionally attending to BCI targets. In addition, the correlation-based safety margin was significantly larger during idle periods, showing that the strongest decoder evidence remained farther from the decision threshold. This separation is also visible in the momentum dynamics: whereas task periods showed progressive evidence accumulation toward a target decision, idle-period momentum rapidly decayed toward the lower bound and remained at a stable plateau, well below the firing threshold (see Fig. 3). Together, these results indicate that the absence of false activations was not merely an isolated behavioral observation, but was consistent with reduced stimulus-locked EEG responses and decoder dynamics that remained separated from the decision boundary.

Because asynchronous visual-BCI studies differ widely in task design, number of targets, EEG setup, calibration duration, decision policy, and definition of idle-state performance, direct quantitative comparison across studies is not straightforward. Table 1 therefore provides a contextual overview of representative approaches rather than a rank-based benchmark. Its purpose is to highlight how non-control handling has been addressed in prior work and to situate the present StAR–Burst implementation with respect to calibration burden, task complexity, EEG practicality, and explicit idle-state evaluation.

**Table 1:** Contextual overview of representative asynchronous visual-BCI studies addressing non-control or idle-state handling. The studies differ substantially in task structure, number of targets, visual-evoked-potential paradigm, calibration procedure, decision rule, EEG setup, idle-state definition, and false-activation metric. Values are therefore reported to situate the present work within the literature, rather than to support direct rank-based comparisons. NR = not clearly reported.

| Study | Task / targets | Calibration | Main reported metric | Timing window | Idle / non-control metric |
| --- | --- | --- | --- | --- | --- |
| Pan et al. (2013) [8] | Binary SSVEP brain switch for control vs idle detection | 320 s (32 trials of 10 s) | 86.67% accuracy and 4.17% FPR in the on-line synchronous test | 7.49 correct operations/min in the asynchronous test | 0.98 false operations/min in the asynchronous test |
| Zhang et al. (2015) [9] | 3-target SSVEP BCI with several idle-state conditions | ~6.3 min | 88.0 ± 11.1% offline control-state accuracy | 3 s EEG epochs | Offline idle FPRs from 7.4 ± 5.6% to 14.2 ± 10.1%; online simulation FPRs as low as ~0.5 event/min in eyes-open idle and 0.05 event/min in eyes-closed idle |
| Du et al. (2019) [10] | 8-target SSVEP work/idle-state detection with multiple idle-state conditions | NR | Work/idle AUC = 0.89 with 1 s data; frequency-recognition accuracy = 96.45 ± 0.14% | 1 s data length | Offline work/idle detection reported mainly as AUC; no online false-activation rate foregrounded |
| Ke et al. (2023) [11] | 14-target asynchronous SSVEP selection with frequency-specific control-state detection | Offline and online training blocks | TPR 93.16 ± 4.4%; balanced accuracy 92.89 ± 4.02%; ITR 124.95 ± 12.35 bits/min | 591.63 ± 5.65 ms average data length | FPR 5.21 ± 5.85% |
| Meng et al. (2025) [12] | Model-based SSVEP brain switch with application demonstrations | Calibration-free for most participants | Reliable switch-gated control; ~0.1 FP/h or better | 9.3–11.2 s triggering time across experiments | No false triggers during the 10-h video-2 experiment |
| Nagel & Spüler (2019) [13] | 32-target VEP matrix speller with non-control detection | 10 min | 99.3 ± 0.43% accuracy; 122.7 bits/min for the 32-target matrix speller | 2.61 ± 0.78 s average trial duration | 0.075 erroneous classifications/min during non-control state |
| Gembler et al. (2020) [14] | 4-target, 32-target, and dictionary-assisted 32-target asynchronous c-VEP spellers | 1.2 min for 4-target / 6.6 min for 32-target | 94.4%, 95.5%, and 94.0% accuracy for 4-target, 32-target, and dictionary-assisted 32-target spelling | 2.31 s, 3.04 s, and 3.16 s for 4-target, 32-target, and dictionary-assisted 32-target spelling | Asynchronous threshold-based implementation allowed a non-control state; no dedicated idle false-activation rate was reported |

Table 1 continued
| Study | Task / targets | Calibration | Main reported metric | Timing window / | Idle / non-control metric |
| --- | --- | --- | --- | --- | --- |
| Zheng et al. (2019) [15] | c-VEP brain switch using 7-class pseudorandom codes for control vs idle detection | 50 training trials (~54 s of stimulation; setup in a few minutes) | Binary brain-switch activation task | 1.72 ± 0.38 s average reaction time | No false-positive events over an average idle time of 183.53 ± 119.73 s |
| Zheng et al. (2022) [16] | Binary c-VEP brain switch for control vs idle detection | NR | Binary trigger task | 1.49–1.67 s average reaction time, depending on operating point | 1.46 $\times 10^{-2}$ FP/min at the faster operating point; no false positives at the more conservative operating point |
| Lai et al. (2024) [17] | 4-target asynchronous c-VEP selection with idle-state handling | ~5 min | PPV ≈99.2–99.7% with m-sequence and DIBS | 2.63–2.75 s reaction time | FPR 6.13 $\times 10^{-2}$ min$^{-1}$ with m-sequence and 7.35 $\times 10^{-2}$ min$^{-1}$ with DIBS |
| **Present study** | **11-class StAR-Burst c-VEP keypad with explicit video-viewing idle condition** | **40 s** | **99.8% online accuracy** | **4.3 s mean decoding time** | **0 observed spurious activations during an explicit 2-min video-viewing idle condition** |

Earlier work has shown that robust non-control handling can be achieved through several routes, including explicit idle-state or control-state classifiers [9, 10, 11], pseudo-key or brain-switch formulations [8, 15, 16], and asynchronous confidence- or threshold-based rejection procedures [13, 14, 17]. These approaches have demonstrated strong non-control performance, but many of them rely on explicit control-vs-idle discrimination, threshold optimization, dedicated non-control modeling or training, or binary brain-switch formulations. For example, Meng *et al*. proposed a highly reliable SSVEP-based brain switch using a virtual physical-system model, reporting false-positive rates of approximately 0.1 events per hour or better, with triggering times on the order of 9–13 s across representative video-viewing and application-oriented experiments, depending on the protocol [12]. This work also demonstrated the switch in application-oriented scenarios, including robotic-arm and wheelchair control. However, the non-control solution was formulated primarily as a brain-switch or gating mechanism that activates downstream control, whereas the present study evaluates idle-state behavior within the same evidence-accumulation decoder used for multi-target c-VEP selection. Similarly, c-VEP brain-switch approaches by Zheng *et al*. achieved very low false-positive rates with reaction times around 1.5–1.7 s [15, 16], whereas asynchronous c-VEP spelling systems such as Gembler *et al*. reported accuracies around 94–95% with mean selection times of 2.3–3.2 s [14]. Lai *et al*. further demonstrated low false-positive rates and high positive predictive values in a 4-target asynchronous c-VEP system with reaction times around 2.6–2.8 s after less than 5 min of calibration [17].

The contribution of the present study is therefore not to claim the first demonstration of robust non-control handling, but to show that idle-state behavior can be incorporated simply and efficiently within the same decoder used for multi-target selection. Here, this was achieved using only 33 s of usable calibration data, a lightweight eight-channel semi-dry EEG system, visually comfortable stimuli, 99.8% control accuracy in an 11-class keypad task, and no observed spurious activations during an explicit 2-minute video-viewing idle condition. This combination is noteworthy because it brings together multi-class decoding, short calibration, high accuracy, and explicit idle-state evaluation within a single user-friendly BCI framework. Having established the screen-based performance of the StAR–Burst framework, the next step was to evaluate a proof-of-concept implementation in a more immersive interaction context.

## 3 VR proof of concept

### 3.1 Participants

This part of the experiment involved the same 30 healthy participants as before (10 women; mean age = 28.4 ± 7.7 years).

### 3.2 Shell-game VR BCI

The VR task was implemented in Unreal Engine^1^ using Blueprints to control game logic, animate objects, and transmit flash onsets to the decoding pipeline via LabStreamingLayer (LSL). To reduce unnecessary stimulation and improve ecological validity, stimulation was automatically suspended for objects outside the participant’s current field of view (i.e., outside the view frustum or fully occluded). Although not essential for the present three-target task, this mechanism is intended to support future scenarios with more potential targets than available codes by temporarily deactivating hidden items and reallocating their codes to visible targets.

Participants performed a simplified shell game. Three cubes (“cups”) were aligned horizontally in front of the participant, and a sphere (the “ball”) was briefly revealed before being hidden under one cube. The three cubes were then shuffled by randomly swapping the cubes’ positions and stopped. During the selection phase, the textured stimulus was rendered as a dynamic material mapped onto each moving cube (i.e., the texture remained locked to the object as the viewpoint changed with head motion) and flashed according to the cube’s assigned Burst code. Once the BCI reached a decision, the ball location was revealed and the selected cube turned green if correct and red otherwise (click here to see the [video showcasing the VR condition](https://nextcloud.isae.fr/index.php/s/9Fzin4Lzk5jJb5E)).

The task was performed using a Meta Quest 3 headset fitted with the Elite head strap. This configuration ensured stable headset placement while maintaining clearance over occipital and parieto-occipital electrodes to limit pressure on the sensors and preserve comfort. The scene geometry was set such that the lateral cubes required head rotation to inspect, ensuring that participants actively moved their head to face the targets during the task.

### 3.3 Protocol

The VR experiment was conducted after completion of the screen-based keypad experiment. This order was kept fixed for all participants and the two conditions were therefore not counterbalanced. Importantly, the VR task was not decoded using the classifier trained during the keypad experiment. Instead, a separate VR-specific calibration phase was performed immediately before the shell-game task, and the classifier used for VR online control was trained only on this VR calibration data.

During this calibration phase, participants focused on a single cube onto which the stimulus texture was overlaid and flickered according to a random Burst code. The calibration phase lasted 43 *s* in total and consisted of five blocks, each composed of three repetitions of a 2.2 *s* code followed by a 2 *s* pause, yielding 33*s* of usable calibration data. Participants then completed 10 trials of the shell-game task. In each trial, they attempted to identify the cube under which the ball was hidden by focusing on the cube they selected. Finally, as in the screen-based condition, participants completed the same subjective ratings assessing visual comfort, peripheral distraction, and mental fatigue, as well as the CVS questionnaire.

### 3.4 Measures

Objective BCI performance was evaluated using the same accuracy and decoding-time metrics as in the keypad experiment. We also examined whether differences in decoding performance were accompanied by differences in stimulus-locked VEP responses. All EEG analyses were performed using the same preprocessing and epoching procedure as described in Section II-H for both keypad and VR data. For each participant, stimulus-locked VEP responses were averaged across trials for both the keypad and VR tasks. From the resulting waveforms, we extracted the mean amplitude, standard deviation of the amplitude, and peak latency within the 150–210 ms time window corresponding to the N2 component, and within the 250–310 ms time window corresponding to the P3 component. These features were extracted at electrode Oz, where the VEP response was most pronounced. For the planned comparisons of N2 and P3 mean amplitudes between keypad and VR conditions, Bonferroni correction was applied, with the significance threshold set to *p* = .025.

Subjective assessment was collected through the CVS questionnaire at the end of the VR task (see section 2.7 for more details).

### 3.5 Results

Normality of the data was assessed using the Shapiro-Wilk test with alpha = 0.05.

#### 3.5.1 Subjective results

Subjective ratings remained consistently favorable as participants reported high comfort (median = 8.0, IQR = 1.0), good peripheral transparency (median = 7.0, IQR = 2.0), and low mental fatigue (median = 2.0, IQR = 2.0). Since the subjective data originate on Likert scales, as ordinal data, they do not adhere to normality assumption. Wilcoxon signed-rank tests revealed a significant difference in mental fatigue between conditions, with participants reporting lower mental fatigue during the Screen task than during the VR task (*z* = *−*2.49, *p* = .012). No significant differences were found for visual comfort (*z* = *−*0.51, *p* = .603) or peripheral transparency (*z* = *−*0.56, *p* = .569).

#### 3.5.2 BCI accuracy and decoding time

Online control in the VR reached 100% accuracy.

Shapiro-Wilk tests showed that neither participant-wise decoding time on the Screen task (*W* = 0.741, *p <* .01) nor decoding time on the VR task (*W* = 0.879, *p* = .03) were normally distributed. Based on this, a Wilcoxon signed-rank test was used, and the median and interquartile range were used to summarize both variables.

A Wilcoxon signed-rank test indicated that decoding times were significantly longer during the VR task (median = 5.4, IQR = 1.0) than during the Screen task (median = 4.3, IQR = 0.9), *z* = 3.59, *p <* .001, *r*_*rb*_ = 0.75.

#### 3.5.3 VEP analysis

At electrode Oz, the keypad condition elicited a significantly larger N2 response, corresponding to a more negative mean amplitude, than the VR condition within the 150–210 ms time window (paired *t*-test, *t*(29) = *−*2.76, *p* = .005, *d*_*z*_ = *−*0.503). Similarly, within the 250–310 ms time window, the P3 component showed a significantly greater positive mean amplitude in the keypad condition relative to the VR condition (paired *t*-test, *t*(29) = 3.8, *p <* .001, *d*_*z*_ = 0.70).

### 3.6 Discussion

In the immersive VR shell-game task, the system likewise achieved 100% accuracy, supporting the feasibility of visually comfortable c-VEP control within a VR headset. However, mean decoding time was significantly increased relative to the screen-based keypad task (5.6 s vs. 4.3 s, *p <* .001), despite the VR task involving only three possible targets. This slower decision-making suggests that the effective evidence accumulation rate was reduced in VR. One potential explanation is that evoked-response amplitudes during calibration appeared smaller in VR than on the monitor (see Fig. 6), which may have reduced the reliability of the inferred 0/1 stream and therefore requires longer accumulation to reach decision thresholds. Second, the headset may have introduced additional constraints on EEG acquisition: pressure from the VR headset straps and repeated left–right head movements during the task can perturb electrode contact and increase motion-related artifacts affecting on-line Burst code reconstruction. These integration challenges are consistent with broader observations in VR–EEG systems, where mechanical constraints and motion can impact signal quality and electrode placement [36]. Future work should therefore prioritize tighter mechanical integration of the VR headset and EEG (e.g., sensor placement and cabling compatible with the headset strap geometry), as well as motion-aware artifact mitigation.

Although mean subjective ratings tended to be slightly lower in VR than on the monitor, paired statistical comparisons revealed no significant differences between conditions for comfort or peripheral transparency. This pattern suggests that the proposed stimulation remained well tolerated when transferred from a standard display to a VR headset. The small numerical trend toward lower ratings in VR is nevertheless plausible, as near-eye displays can impose additional visual and oculomotor demands associated with digital eye strain and discomfort in VR [37, 38].

Participants did report significantly higher mental fatigue in the VR condition. However, this result should be interpreted cautiously, because the monitor-based keypad task was always completed before the VR task and the two conditions were not counterbalanced in this proof-of-concept study. The higher mental fatigue ratings in VR may therefore reflect not only the immersive display itself, but also an order effect due to accumulated fatigue over time. Overall, these findings suggest that comfort-optimized stimulation generalizes well to immersive VR, while motivating further refinements of rendering parameters, such as luminance, contrast, and texture-amplitude depth, as well as headset fit for longer sessions.

## 4 General conclusion

We demonstrated a fast-calibrated, visually comfortable c-VEP rBCI that achieved 99.8% online accuracy in a screen-based 11-class keypad task with only 33 s of calibration data, and 100% accuracy in an immersive VR shell-game task, using a practical *semi-dry* EEG system. Compared with our earlier dry-electrode implementations [20, 19], the present semi-dry setup was associated with strong evoked responses and highly reliable code reconstruction, although no direct dry-versus-semi-dry comparison was performed. During the explicit video-viewing idle condition, no spurious activations were observed, and complementary SNR and safety-margin analyses indicated that idle-state activity remained clearly separated from intentional control under the tested conditions.

While VR control remained perfectly accurate, decisions were significantly slower in VR in this fixed-order proof-of-concept design, consistent with possible motion- and headset-related constraints, although this interpretation is limited by the fixed task order and the absence of counterbalancing. Overall, these findings support the feasibility of extending comfort-optimized c-VEP interaction beyond standard laboratory settings and into immersive VR, while highlighting the importance of improved EEG–VR headset integration for faster and more robust embodied interaction.

Despite this progress, the study has several limitations. First, the experimental conditions were not counterbalanced, which may have introduced order effects, particularly for subjective fatigue ratings. Second, the idle-state evaluation was limited to a single 2-minute video-viewing period. Future work should therefore assess idle-state behavior over longer durations and in more dynamic scenarios, including protocols that alternate between idle periods and active commands to test whether the BCI remains both conservative during non-control and responsive after idle intervals. Finally, the present implementation relied on a generic decision threshold that was applied across participants. Further improvements may be achieved through user-adaptive thresholding strategies that account for inter-individual variability in signal quality, decoder confidence, and idle-state activity.

## Ethics statement

The study protocol was approved by the University of Toulouse ethics committee (CER approval number 2025_1012) and complied with the Declaration of Helsinki. All participants provided written informed consent prior to participation.

## Data and code availability

Data and code availability information should be completed by the authors before posting the preprint, if a public repository or access procedure is available.

## Author contributions

Author-contribution information should be completed by the authors before posting the preprint.

## Acknowledgements

Acknowledgements, if applicable, should be added here.

## Footnotes

1 https://www.unrealengine.com

